# Risk Assessment of the newly emerged H7N9 avian influenza viruses

**DOI:** 10.1101/2022.11.11.516200

**Authors:** Pengxiang Chang, Jean-Remy Sadeyen, Sushant Bhat, Rebecca Daines, Altaf Hussain, Huseyin Yilmaz, Munir Iqbal

## Abstract

Since the first human case in 2013, H7N9 avian influenza viruses (AIVs) have caused more than 1500 human infections with a mortality rate of approximately 40%. Despite large-scale poultry vaccination regimes across China, the H7N9 AIVs continue to persist and evolve rapidly in poultry. Recently, several strains of H7N9 AIVs have been isolated and shown the ability to escape vaccine-induced immunity. To assess the zoonotic risk of the recent H7N9 AIV isolates, we rescued viruses with hemagglutinin (HA) and neuraminidase (NA) from these H7N9 AIVs and six internal segments from PR8 virus (A/Puerto Rico/8/34 [H1N1]) and characterized their receptor binding, pH of fusion, thermal stability, plaque morphology and in ovo virus replication. We also assessed the cross-reactivity of the viruses with human monoclonal antibodies (mAbs) against H7N9 HA and ferret antisera against H7N9 AIV candidate vaccines. The H7N9 AIVs from the early epidemic waves had dual sialic acid receptor binding characteristics, whereas the more recent H7N9 AIVs completely lost or retained only weak human sialic acid receptor binding. Compared with the H7N9 AIVs from early epidemic waves (2013-2016), the recent (2020/21) viruses formed larger plaques and increased replication titres in ovo, demonstrating increased acid stability but reduced thermal stability. Further analysis showed that these recent H7N9 AIVs had poor cross-reactivity with the human mAbs and ferret antisera, highlighting the need to update the vaccine candidates. To conclude, the newly emerged H7N9 AIVs showed characteristics of typical AIVs, posing reduced zoonotic risk but a heightened threat for poultry.

## INTRODUCTION

A new H7N9 virus strain emerged in China in 2013 and has resulted in more than 1500 confirmed human infections with ∼40% case fatality rate (1). During the first four epidemic waves (2013-2016), only the low-pathogenicity avian influenza (LPAI) genotype of H7N9 avian influenza viruses (AIVs) were prevalent, however, the high-pathogenicity avian influenza virus (HPAI) genotype emerged in the fifth epidemic wave (2016-2017) and has since been co-circulating with the LPAI H7N9 AIVs in China (2). Compared to the LPAI, the HPAI H7N9 phenotype can cause more severe clinical diseases in chicken, ducks, mice as well as in ferrets (3-5). In the early epidemic waves, the H7N9 AIVs were mainly reported near the Yangtze delta region and Pearl delta region in China. Surprisingly, the H7N9 AIVs spread geographically all over China during fifth epidemic wave, with 33 out of 34 provinces or municipalities in mainland China affected (6). Furthermore, possible hospital and family clusters of H7N9 AIV have been reported and raised concern of human-to-human transmission and thus possible pandemic potential (7). Given the threat posed by H7N9 AIVs to both human and animal health, the Chinese government implemented a mass vaccination program in poultry against H7N9 from September 2017. Since then, human H7N9 AIV infection dropped sharply, with no new cases reported since 2020. The successful prevention of human infection with H7N9 AIVs by large scale vaccination of poultry relies on the following two factors: On one hand, the mass poultry vaccination substantially reduces the prevalence of H7N9 AIVs in poultry, consequently reducing the risk of human infections as the majority of human H7N9 cases are linked to exposure to infected poultry or contaminated environments. On the other hand, when some viruses escape the vaccine-induced immunity, they pose a lower zoonotic risk to humans as the vaccine escape mutations resulted in a loss or weakened affinity to human receptor binding (8, 9).

Recently, Chen *et al*. (10) isolated several H7N9 AIVs from poultry about six months post-vaccination with H7N9 H7-Re3 and H7N9 rLN79 vaccine strains in China. Compared with the vaccine strains, the recently emerged H7N9 AIVs showed an 8-to-128-fold reduction in the antigenic cross-reactivity with chicken post-vaccination sera as measured by hemagglutinin inhibition (HI) assay. Of note, the H7N9 A/Chicken/Yunnan/1004/2021 could evade the vaccine protection and caused 40%−80% mortality in immunized chickens. Due to the antigenic drift, these newly emerged H7N9 AIVs might gradually become dominant under the vaccine-induced pressure. It is therefore important to closely monitor the evolution and thus the zoonotic risk of these newly emerged H7N9 AIVs.

Studies investigating transmission of H5N1 AIVs in ferret model revealed that efficient human-to-human transmission requires shift from avian to human receptor binding preferences, reduced pH of fusion and increased HA thermal stability (11, 12). The receptor binding specificity of influenza A virus is one of the major determinants of host specificity; both H2N2 and H3N2 pandemics have been linked to the preference switch from avian (α2,3-linked sialic acid) to human (α2,6-linked sialic acid) receptor specificity (13). The H7N9 AIV isolates from the early epidemic waves showed comparable binding preferences to both receptors, a characteristic of H7N9 AIV which has been associated with the zoonotic infections of humans in China since 2013 (14). The acid and thermal stability of HA have also been shown to play important role in both interspecies and intraspecies transmission of influenza viruses (reviewed in [15]). Generally, the fusion pH of AIVs ranges from 5.6 to 6.2, while human influenza viruses require relatively lower pH (5.0 to 5.4) for efficient fusion (16). The H7N9 AIV from the early epidemic waves showed a pH of fusion of around 5.6 to 5.8. However, some HPAI H7N9 in the fifth epidemic wave appears to fuse at pH 5.4 or lower (17, 18). In terms of HA thermal stability, the H7N9 AIVs appeared progressively more thermal stably, the LPAI and HPAI H7N9 AIVs from epidemic wave five have been shown to be comparable to or more thermal stable than the 2009 H1N1 pandemic virus (19).

In this study, we systematically investigated the risk of the newly emerged H7N9 AIVs to human and animal health by assessing the virus receptor binding, pH of fusion, HA thermal stability, virus infectivity and the cross-reactivity of these H7N9 variants with human H7N9 mAbs and ferret sera raised against World Health Organization (WHO) vaccine candidates.

## Results

### Newly emerged H7N9 AIVs lost or retained weak human-like sialic acid receptor binding

The receptor binding switch from avian-like to human-like is the key prerequisite for the interspecies transmission of AIVs from birds to humans. To examine such properties of the H7N9 AIVs in China, we utilized biolayer interferometry to characterize the receptor-binding profiles of these viruses to the avian-like 3′-sialylacetyllactosamine (3SLN) and human-like 6′-sialylacetyllactosamine (6SLN) receptor analogues. Three H7N9 AIVs isolated from December 2020 to March 2021 from three different provinces in China were selected for this study; they are HPAI H7N9 A/Chicken/Yunnan/1004/2021 (referred to as YN/21) from Yunnan province, south of China, HPAI H7N9 A/Chicken/Hebei/1009/2020 (referred to as HB/20) from Hubei Province, north of China, and HPAI H7N9 A/Chicken/Shanxi/1012/2021 (referred to as SX/21) from Shanxi province, north of China (10). All viruses were isolated after mass vaccination with H7N9 H7-Re3 (Harbin Weike Biotechnology Development Company) or rLN79 (Guangzhou South China Biologic Medicine) vaccines and showed reduced antigenic cross-reactivity with the vaccine strains. We also included H7N9 A/Anhui/1/2013 from the first epidemic wave (referred to as AH/13), and HPAI H7N9 AIV A/Guangdong/17SF003/2016 (referred to as GD/16) from epidemic wave five for reference. The recombinant viruses were generated by reverse genetics with HA and NA from these H7N9 AIVs and six internal segments from PR8 virus (A/Puerto Rico/8/34 [H1N1]). In agreement with our previous reports (9, 20), the H7N9 AH/13 showed comparable binding to both 3SLN and 6SLN receptor analogues **(Figure 1)**. In comparison to the AH/13, the GD/16 from the epidemic wave five showed significantly stronger binding to 3SLN and a slight increase in binding to 6SLN receptor analogues respectively (∼293-fold increase to 3SLN and ∼2-fold increase to 6SLN). The HB/20 and SX/21 HPAI H7N9 AIVs completely lost binding to the 6SLN receptor analogue while gaining a slight improvement of binding to the 3SLN receptor analogue when compared with AH/13 (∼4-fold and 2-fold increase to 3SLN respectively). YN/21 showed similar binding affinity toward 3SLN receptor analogue as HB/20 and SX/21. However, it still bound to 6SLN analogue, though the binding affinity was ∼39 folds weaker when compared with the AH/13. In conclusion, the newly emerged H7N9 AIVs showed typical receptor binding profiles of AIVs with complete loss or very weak binding toward human-like receptors.

**FIG 1.**
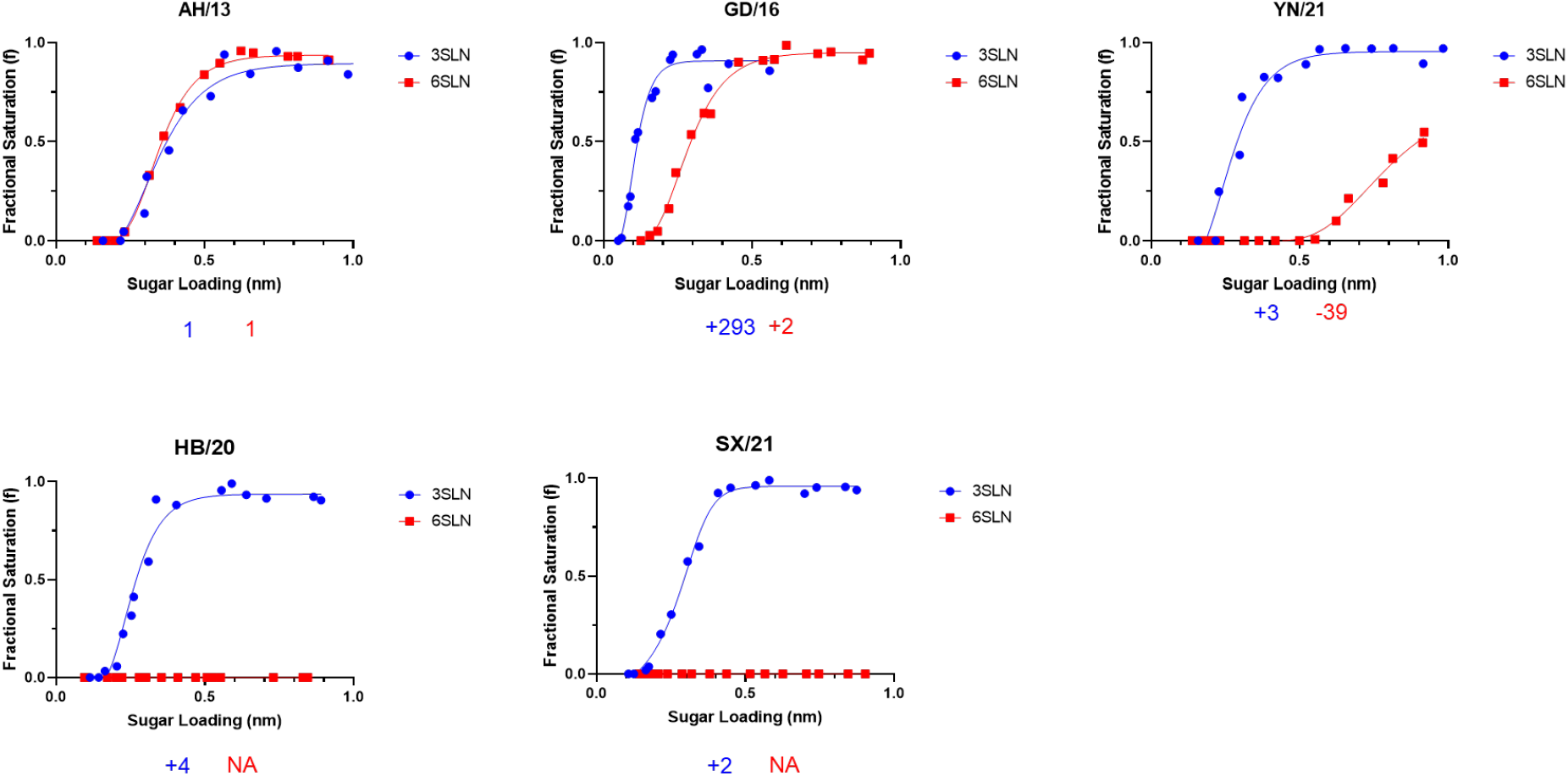
Receptor binding profiles of newly emerged H7N9 AIVs. The binding of purified recombinant H7N9 AIVs to avian and human receptor analogues was measured by biolayer interferometry. The H7N9 AIVs was generated by reverse genetics with HA and NA from indicated H7N9 AIVs and the internal segments from PR8 H1N1 virus The numbers below each figure show the fold change of receptor binding of indicated viruses to avian (α2,3-SLN, shown in blue) and human (α2,6-SLN, shown in red) receptor analogues as compared to recombinant AH/13. ‘-’ indicates reduction, ‘+’ indicates increase and ‘NA’ = not applicable. Data is the combination of two repeats for each virus and receptor analogue combination.

### Newly emerged H7N9 AIVs demonstrated reduced pH of fusion and HA thermal stability

The pH of fusion has been shown to play an important role in host adaptation and transmission; human-adapted influenza viruses generally fuse at a lower pH, while AIVs normally have a higher pH fusion threshold (16). Consistent with the previous report, AH/13 triggered fusion at pH 5.6 (21). Surprisingly, the GD/16 from the epidemic wave five fused at pH 5.3 or lower **(Figure 2A)**. Compared with the AH/13, all the newly emerged H7N9 AIVs showed a lower threshold pH for HA activation, with YN/21 fused at pH 5.4 or lower and HB/20 and SX/21 fused at pH 5.5 or lower. In addition to receptor binding and pH of fusion, increased HA thermal stability has been linked to a vital role in AIV evolution (22). Previous work has shown that the HA of aerosol or respiratory droplet-transmissible influenza viruses are comparatively more thermal stable (11, 12). To test the heat stability of the HA protein, the newly emerged recombinant H7N9 AIVs together with GD/16 and AH/13 viruses were incubated at 4.0°C, 50°C, 50.7°C, 51.9°C, 53.8°C, 56.1°C, 58.0°C, 59.2°C and 60°C for 30 min, after which the loss of HA activity was determined. Compared with the AH/13 from the epidemic wave one, the GD/16 showed increased thermal stability **(Figure 2B)**. All the newly emerged H7N9 AIVs were sensitive to high temperature, they lost HA activity more rapidly than both AH/13 and GD/16.

**FIG 2.**
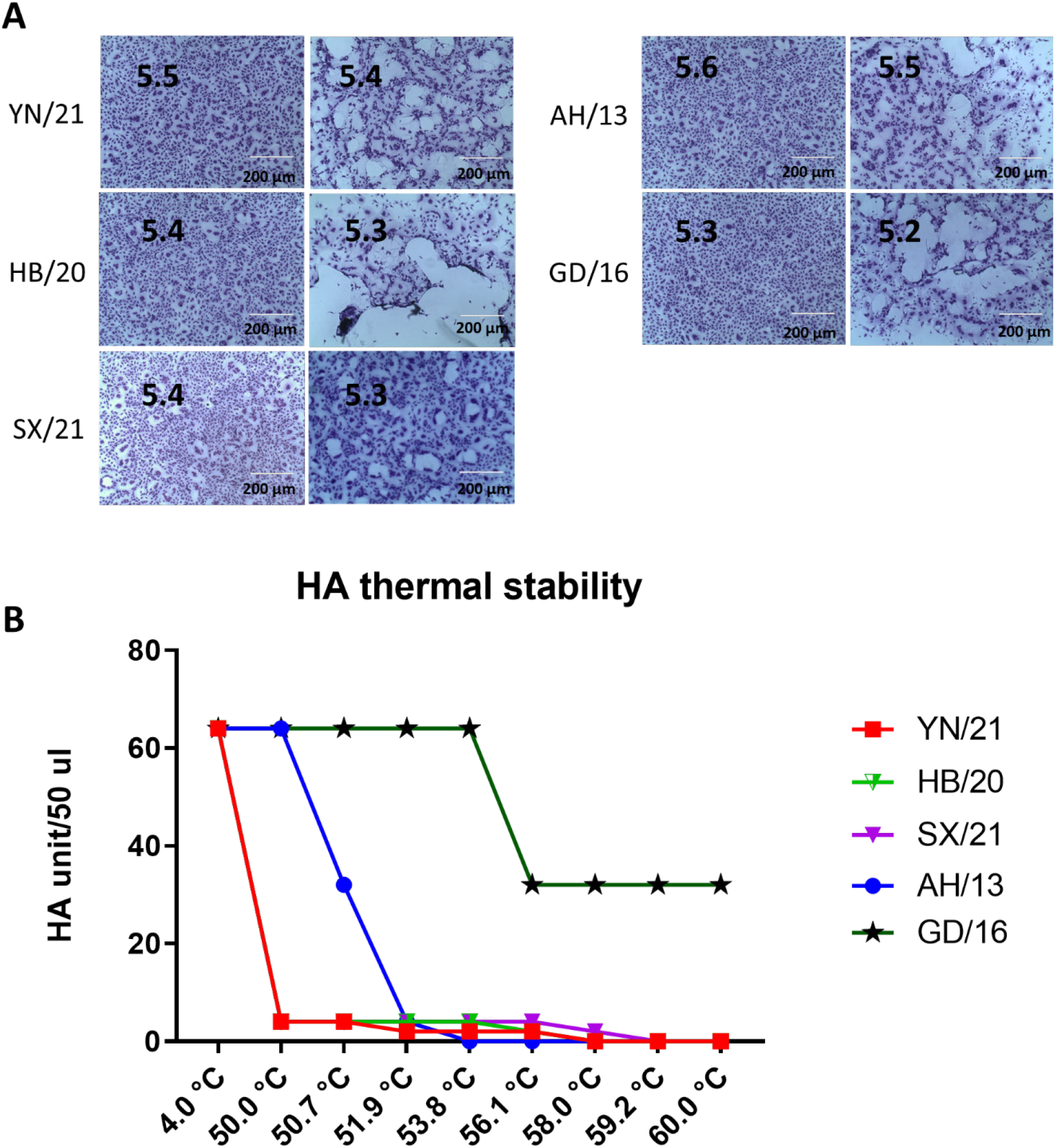
pH fusion and thermal stability of newly emerged H7N9 AIVs. **(A)** Syncytium formation in Vero cells infected with recombinant H7N9 AIVs (HB/20, SX/21 and YN/21), GD/16 and AH/13 with the HA and NA from H7N9 AIVs and the internal segments from PR8 H1N1 virus. The pH, at which 50% of maximum syncytium formation was observed, was taken as the predicted pH of fusion, shown on the left panel. The syncytium formation at 0.1 pH unit lower than the fusion threshold were shown on the right panel as control. Results shown are representative of three experimental repeats. **(B)** HA thermal stability of recombinant H7N9 AIVs. 64 HA units of recombinant virus were either left at 4.0°C as control or heated at 50°C, 50.7°C, 51.9°C, 53.8°C, 56.1°C, 58.0°C, 59.2°C and 60°C for 30 min before the HA assay. Results shown are representative of three experimental repeats.

### The newly emerged H7N9 AIV formed larger plaques and replicated robustly *in ovo*

To assess the fitness of the virus, we assessed the plaque sizes and the virus replication in the embryonated eggs. Alike the Anhui/13 virus, the GD/16 formed relatively small size plaques in Madin-Darby canine kidney (MDCK) cells. However, all the newly emerged H7N9 AIVs formed significantly larger size plaques than the viruses from the early epidemic waves **(Figure 3A and Figure 3B)**. To further assess the effects of these mutations on virus replication, 10-day old embryonated eggs were inoculated with 100 plaque-forming units of the H7N9 AIVs for 48 hours and virus titres were determined by HA assay **(Figure 3C)**. In agreement with the plaque morphology, all the newly emerged H7N9 AIV grew to significant higher titres than the AH/13 and GD/16. To conclude, the newly emerged H7N9 AIV formed larger plaques and replicated robustly *in ovo*, indicating that these newly emerged H7N9 AIVs have more replication fitness than the viruses from the early epidemic waves.

**FIG 3.**
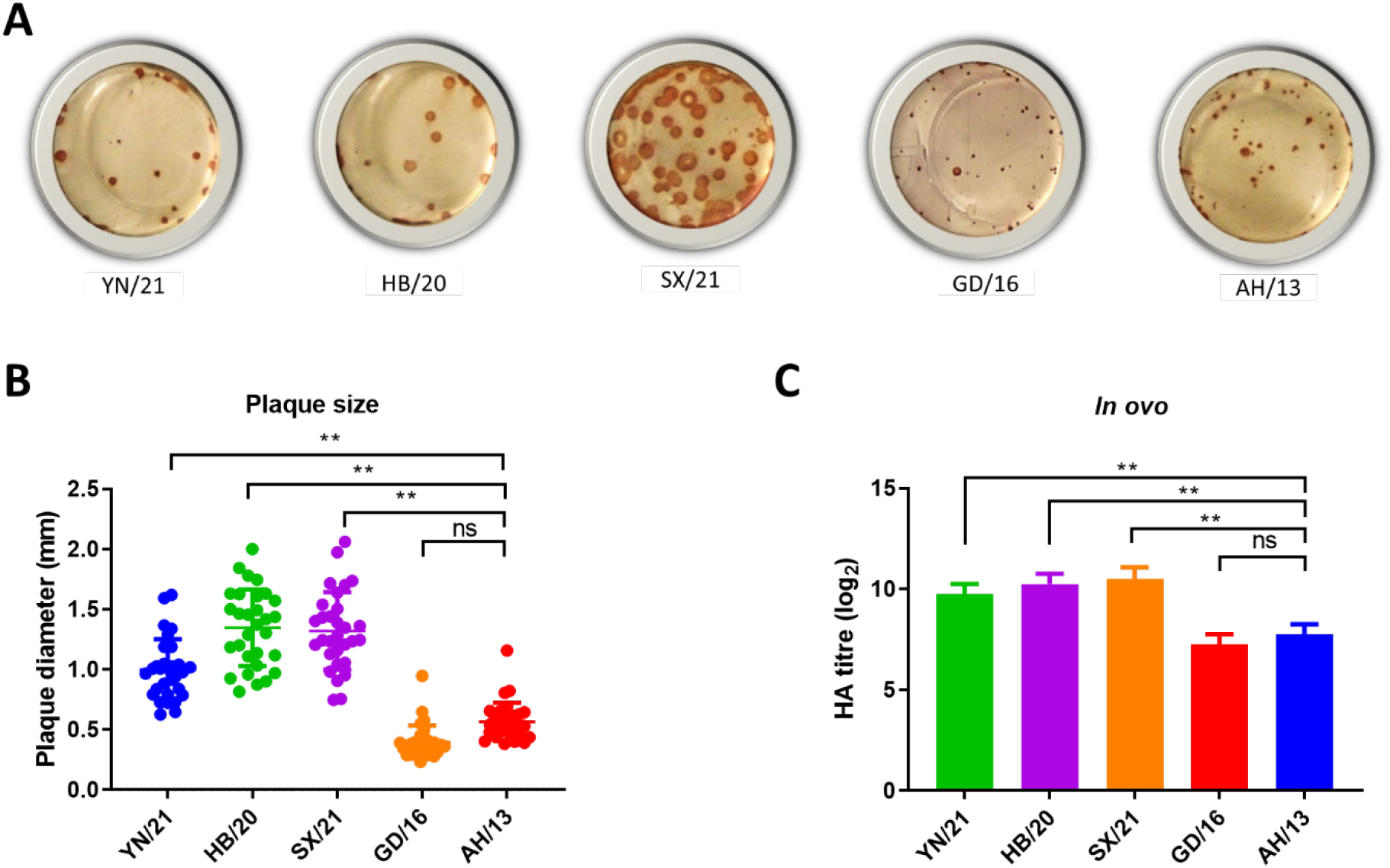
Plaque morphology and virus replication of the newly emerged H7N9 AIVs in MDCK cells and embryonated chicken eggs, respectively. **(A)** The plaque assay was performed in MDCK cells with recombinant newly emerged H7N9 AIVs (HB/20, SX/21 and YN/21), GD/16 and AH/13 with the HA and NA from H7N9 AIVs and the internal segments from PR8 H1N1 virus. **(B)** The plaque size for the recombinant newly emerged H7N9 AIVs (HB/20, SX/21 and YN/21), GD/16 and AH/13 in MDCK cells was measured by selecting 30 random plaques and plaque diameter was measured **(C)** The 10-day old embryonated eggs were inoculated with 100 plaque-forming units of the H7N9 AIVs for 48 h before allantoic fluid was harvested and the viral titres determined by HA assay. Error bar = standard deviation. *p < 0.05; **p < 0.001; ‘ns’ = not significant. Results shown are representative of two experimental repeats.

### The newly emerged H7N9 AIVs reduced/lost cross-reactivity with human monoclonal antibodies

We previously isolated and generated four monoclonal antibodies (mAbs) from humans naturally infected with H7N9 A/Anhui/1/2013 (20, 23). The mAb L3A-44, K9B-122, L4A-14 and L4B-18 target HA residues 125 (Alanine (A)), 133 (Glycine (G)), 149 (Asparagine (N)), and 217 (Leucine (L)), respectively. To investigate the cross-reactivity of the newly emerged H7N9 AIVs with these mAbs, all the newly emerged H7N9 AIVs were tested by HI assay with the panel of mAbs with AH/13 and GD/16 as reference. In line with our previous reports, all four human mAbs showed high HI titres to AH/13 **(Table 1)**. The HI titres of L3A-44, L4B-18 and K9B-122 dropped dramatically while L4A-14 only showed one log2 reduction to GD/16 when compared with AH/13. All the newly emerged H7N9 AIVs completely lost their cross-reactivity with human mAb L4A-14, L3A-44 and K9B-122, with only weak cross-reactivity found with L4B-18. To investigate the cause for the reduced cross-reactivity between mAbs and the newly emerged H7N9 AIVs, the HA amino acid sequences of these H7N9 AIVs were analysed. All the newly emerged H7N9 AIVs harbour Threonine (T) at amino acid residue 125 and 151 **(Table 2)**, which introduced an N-linked glycosylation at amino acid residues 123 and 149 (9); such introduction has been shown to abolish the binding to L3A-44 and L4A-14 respectively (20). The drop in HI titre of L4B-18 with the newly emerged H7N9 AIVs might be due to the L217Q substitution in HA, which has been shown to reduce its cross-reactivity with L4B-18. As for K9B-122, it lost cross-reactivity in the presence of T at HA amino acid residue 123 or Q at HA amino acid residue 217 from all the newly emerged H7N9 AIVs.

**Table 1.**
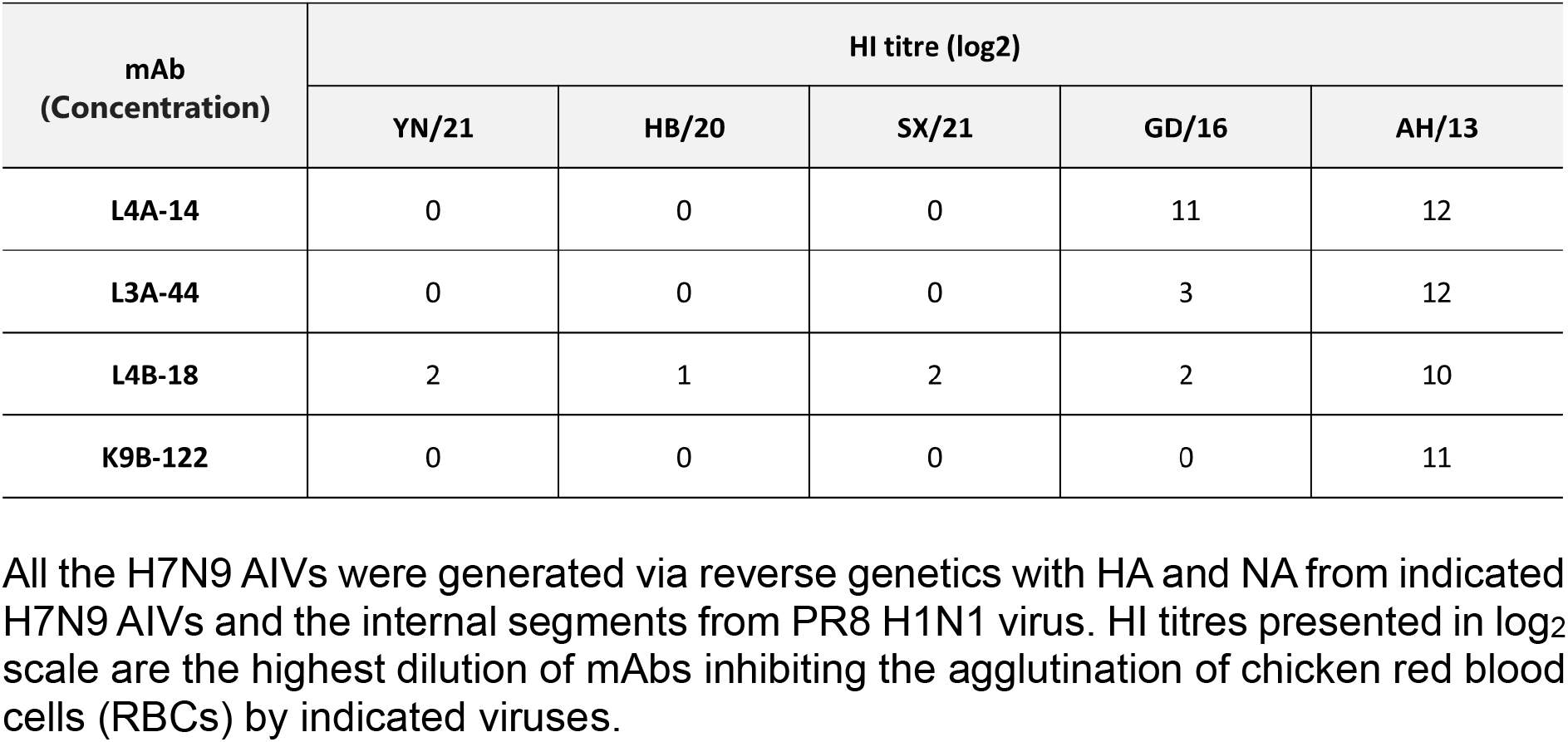
Cross-reactivity of mAbs with the newly emerged H7N9 AIVs.

| mAb<br>(Concentration) | HI titre (log2) |  |  |  |  |
| --- | --- | --- | --- | --- | --- |
|  | YN/21 | HB/20 | SX/21 | GD/16 | AH/13 |
| L4A-14 | 0 | 0 | 0 | 11 | 12 |
| L3A-44 | 0 | 0 | 0 | 3 | 12 |
| L4B-18 | 2 | 1 | 2 | 2 | 10 |
| K9B-122 | 0 | 0 | 0 | 0 | 11 |

All the H7N9 AIVs were generated via reverse genetics with HA and NA from indicated H7N9 AIVs and the internal segments from PR8 H1N1 virus. HI titres presented in log<sub>2</sub> scale are the highest dilution of mAbs inhibiting the agglutination of chicken red blood cells (RBCs) by indicated viruses.

**Table 2.**
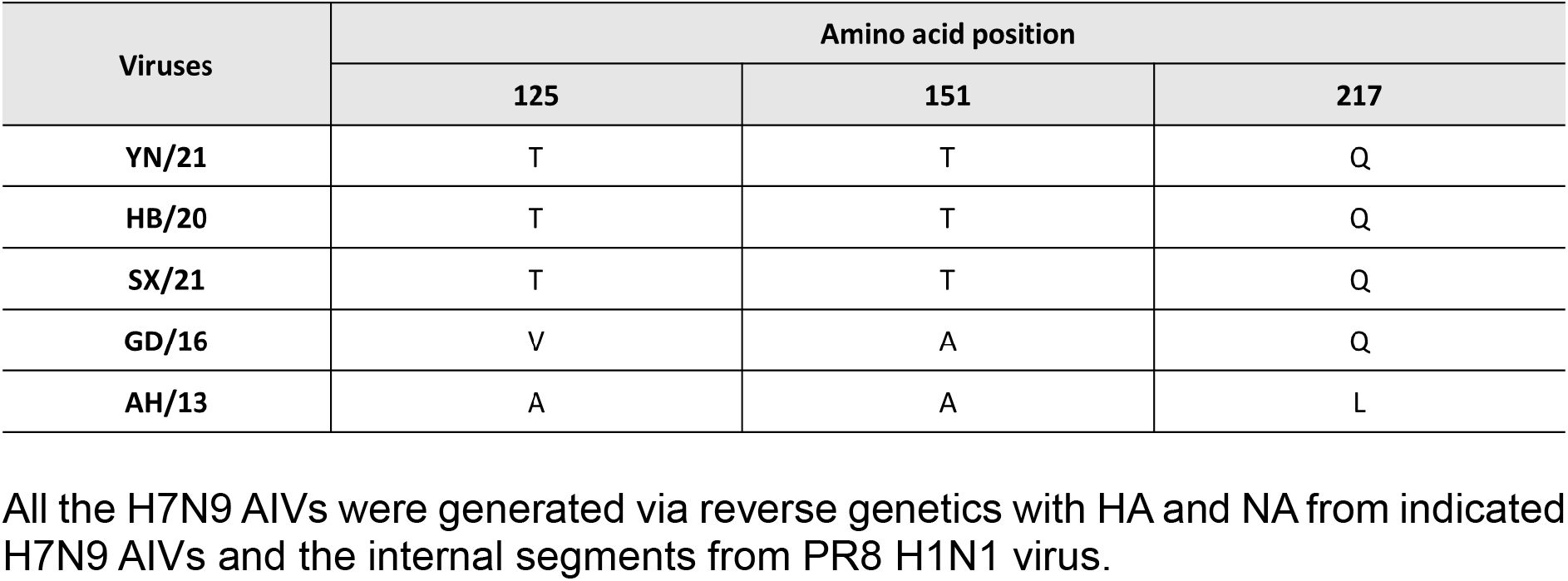
Amino acid residues of newly emerged H7N9 AIV HAs at position 125, 151, and 217.

**Table 3.** Cross-reactivity of newly emerged H7N9 AIVs with sera against WHO vaccine candidates.

| Viruses | HI titre (log2) |  |
| --- | --- | --- |
|  | AH/13 ferret serum | GD/16 ferret serum |
| AH/13 | 9 | 10 |
| GD/16 | 4 | 7 |
| YN/21 | 5 | 5 |
| HB/20 | 5 | 6 |
| SX/21 | 7 | 8 |

All the H7N9 AIV was generated via reverse genetics with HA and NA from indicated H7N9 AIVs and the internal segments from PR8 H1N1 virus. HI titres presented in log<sub>2</sub> scale are the highest dilution of ferret serum inhibiting the agglutination of RBCs by indicated viruses.

### Cross-reactivity of newly emerged H7N9 AIVs with the ferret antisera against H7N9 vaccine candidates

Antigenic characterization of AIVs by ferret antisera has and continues to be used by the WHO for the selection of the candidate vaccine viruses (CVVs) for influenza (24). To test whether the CVVs are still effective against the newly emerged H7N9 AIVs, we assessed the neutralizing profile of ferret antisera against candidate vaccine virus AH/13 from epidemic wave one and GD/16 from epidemic wave five by both HI an microneutralization (MN) assay. Consistent with our previous report (25), GD/16 showed a 5 log_2_ reduction in HI titre with AH/13 ferret antiserum when compared with the AH/13 **(Table 1)**. Likewise, the HI titre of newly emerged H7N9 AIVs YN/21, HB/20 and SX/21 also dropped significantly, except for SX/21 which displayed only a 2 log2 reduction in HI titre. As for the GD/16 ferret serum, surprisingly, the AH/13 showed 3 log_2_ higher HI titre than the GD/16. No significant drop in cross-reactivity was observed between the newly emerged H7N9 AIVs and the GD/16; only YN/21 showed 2 log_2_ reduction in HI titre when compared with the GD/16. Consistent with the HI assay, the MN titre of AH/13 ferret serum to the newly emerged H7N9 AIVs and GD/16 dropped dramatically to below detection level. As to the GD/16 ferret serum, no significant difference in MN titre was observed between AH/13 and GD/16, despite the 3 log_2_ higher HI titre found with AH/13 **(Figure 4B)**. Though only a minor difference in HI titre was observed between GD/16 and the newly emerged H7N9 AIVs, the MN titre to the newly emerged H7N9 AIVs dropped dramatically compared with the GD/16. To conclude, the newly emerged H7N9 AIVs showed poor cross-reactivity with ferret sera raised against the WHO vaccine candidate AH/13 and GD/16, and there was discrepancy in antigenicity measured by HI and MN assay

**FIG 4.**
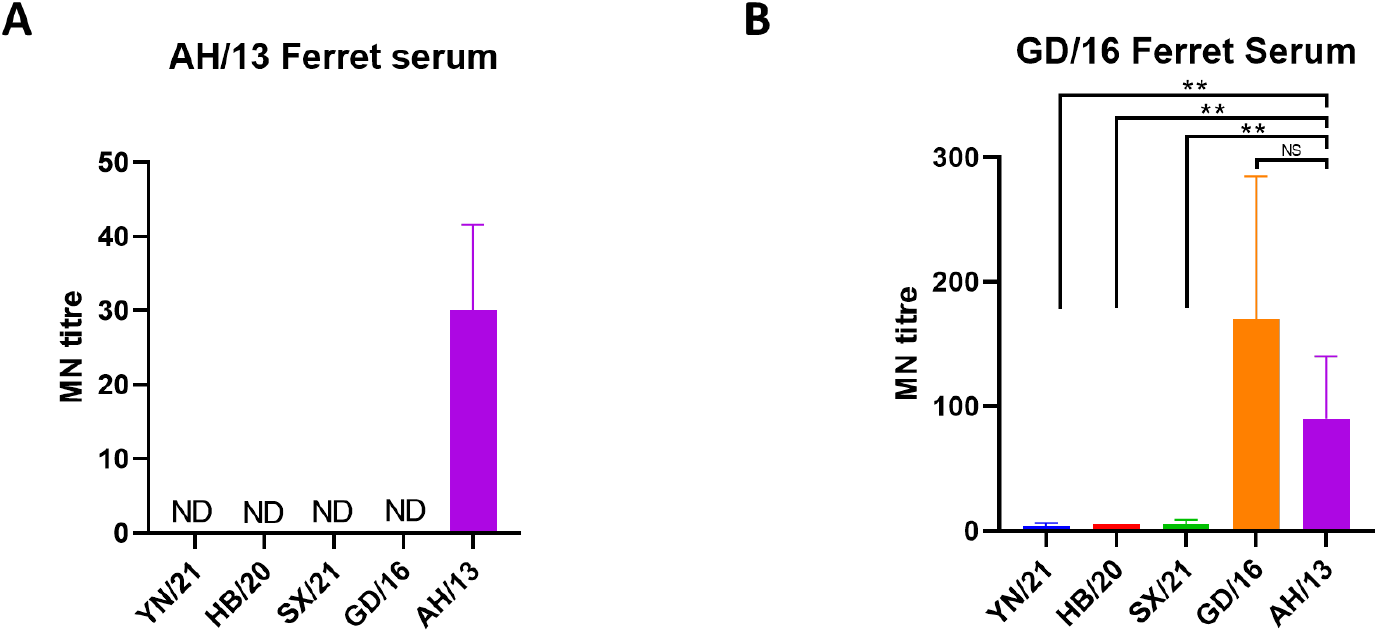
Neutralization of newly emerged H7N9 AIVs by (A) ferret antisera raised against AH/13 and (B) GD/16. The microneutralization assay was performed in MDCK cells with recombinant newly emerged H7N9 AIVs: HB/20, SX/21, YN/21, GD/16 and AH/13 with the HA and NA from H7N9 AIVs and the internal segments from PR8 H1N1 virus. The virus neutralisation titre was expressed as the reciprocal of the highest serum dilution at which virus infection is blocked and the cells survive. Data are presented as mean ± SD and analysed by one-way ANOVA followed by Tukey’s multiple comparison test. Error bar = standard deviation. * p < 0.05; ‘NS’ = not significant ‘ND’ = no detection. ‘MN’ = microneutralization.

## DISCUSSION

To closely monitor the zoonotic risk of the newly emerged H7N9 AIVs, we selected three newly emerged H7N9 AIVs from different provinces of China from 2020 to 2021. Here we showed that the newly emerged H7N9 AIVs had typical AIV-like receptor binding profiles and reduced thermal stability, therefore posing a reduced zoonotic risk when compared to viruses from early epidemic waves. There was very low cross-reactivity of the newly emerged H7N9 AIV with both human mAbs isolated from humans naturally infected with H7N9 AH/13 and the ferret antisera against WHO candidate vaccines AH/13 and GD/16, highlighting the need to update the candidate vaccine strain. In addition to avian-like receptor binding profile, these newly emerged H7N9 AIVs showed increased replication fitness as evidenced by larger plaque sizes and higher replication titres *in ovo*. Moreover, these viruses also demonstrated reduced pH fusion thresholds, which has been shown to be essential airborne transmissibility for H9N2 and H5N1 AIVs between birds (26, 27). Taking all these factors into account, these newly emerged H7N9 AIVs might have increased fitness in poultry than the isolates from the early waves.

The receptor binding specificity of influenza A virus is one of the major determinants of host specificity. It is a prerequisite for pandemic viruses to switch receptor binding specificity from avian receptor α2,3-SA to human receptor α2,6-SA (13). Here we showed that both AH/13 and GD/16 from early H7N9 AIV epidemic waves showed binding towards both avian-like 3SLN and human-like 6SLN receptor analogues. However, among these newly emerged H7N9 AIVs, HB/20 and SX/21 completely lost the receptor binding towards human-like 6SLN receptor analogues while YN/21 showed ∼39 folds of reduction in its binding towards human-like 6SLN receptor analogues when compared with AH/13. We have previously shown that the threonine at residues 125 and 151 in HA formed N-linked glycosylation motifs (N-X-T/S, where X is any amino acid other than proline) and resulted in the addition of glycan to the asparagine at amino acid residue 123 and 149 (9). The receptor binding assay showed that the double glycosylation at amino acids 123 and 149 of H7N9 HA are responsible for the loss of the human-like receptor binding. Here the amino acid sequence analysis showed that all the newly emerged H7N9 AIVs have threonine at residues 125 and 151 of HA and form the N-linked glycosylation motifs **(Table 2)**. Therefore, the drop in binding affinity toward human-like receptor is most probably due to the double glycosylation as a result of threonine at residues 125 and 151 in HA of all the newly emerged H7N9 AIVs. Therefore, if any amino acid change in the HA of these newly emerged H7N9 AIVs that remove the N-linked glycosylation motifs probably will enable the viruses to regain the dual receptor binding properties. On that account, the amino acid changes around amino acid residue 125 and 151 must be closely monitored for the zoonotic risk assessment.

To conclude, we selected H7N9 AIVs that are antigenic distant to the vaccine strains, which are likely to become dominant in the coming years under vaccine-induced pressure. Moreover, the viruses are from different provinces of China, therefore, the study here should provide a comprehensive risk assessment update about the H7N9 AIVs in China. We demonstrated that they posed reduced zoonotic risk but potential heightened threat for poultry by using both *in vitro* and *in ovo* methods. Given the low risk to human infections, we believe the data here is of great importance for the policymakers to consider whether to keep the current vaccine scheme or adopt measures to eliminate the viruses from poultry. However, it is noteworthy that the work performed in this study comprised of *in vitro* and *in ovo* data with recombinant H7N9 AIVs with six internal gene segments from PR8 H1N1 influenza virus; *in vivo* experimentation and utilizing full wild-type viruses in the respective hosts would greatly strengthen the data generated in this study.

## MATERIALS AND METHODS

### Ethics Statement

All the procedures involving embryonated eggs were carried out in strict accordance with the guidance and regulations of United Kingdom Home Office regulations under project licence number P68D44CF4. As part of this process, the work has undergone scrutiny and approval by the animal welfare ethical review board at the Pirbright Institute. All the influenza virus related work was carried out in biosafety level-2 conditions.

### Viruses and cells

The recombinant H7N9 viruses containing HA and NA from LPAI H7N9 (A/Anhui/1/2013) (referred to as Anhui/13), HA and NA from HPAI H7N9 virus (A/Guangdong/17SF003/2016) (referred to as GD/16), HA and NA from HPAI H7N9 virus (A/Chicken/Yunnan/1004/2021) (referred to as YN/21), HA and NA from HPAI H7N9 virus (A/Chicken/Hebei/1009/2020, referred to as HB/20) or HA and NA from HPAI H7N9 virus (A/Chicken/Shanxi/1012/2021, referred to as SX/21). The DNA sequences for the HA and NA of H7N9 viruses are derived from the Global Initiative on Sharing All Influenza Data (GISAID). The GISAID accession numbers for the HA of Anhui/13, GD/16, YN/21, SX/21 and HB/20 are EPI439507, EPI439509, EPI1856859, EPI1857090 and EPI1857209, respectively. The GISAID accession numbers for the NA of Anhui/13, GD/16, YN/21, SX/21 and HB/20 are EPI919607, EPI1010185, EPI1856861, EPI1857092 and EPI1857211, respectively. The HAs and NAs were synthesized by GeneArt (Thermo Fisher Scientific) and then subcloned into PHW2000 plasmids. The polybasic amino acids motif of the HPAI H7N9 HAs were removed. The internal gene segments from PR8 H1N1 virus (A/Puerto Rico/8/34) were generated as previously described (8). Rescued viruses were propagated in 10-day-old embryonated chicken eggs and virus stocks were kept at −80 °C.

The MDCK cells, human embryonic kidney 293T and Vero cells (ATCC) were maintained with Dulbecco’s Modified Eagle’s medium (DMEM) (Gibco), supplemented with 100 U/mL penicillin, 100 μg/mL streptomycin (Gibco) and 10% fetal calf serum (FCS) (Gibco), at 37 °C with 5% CO_2_.

### Monoclonal antibodies and sera

The monoclonal antibodies L4A-14, K9B-122, L3A-44 and L4B-18 were generated by co-transfection with antibody light and heavy chain plasmids using ExpiFectamine™ CHO Transfection Kit according to the manufacturer’s protocol. The supernatants containing antibodies were filtered and purified by HiTrap Protein G HP antibody purification columns (GE Healthcare).

Post-infection ferret antiserum raised against HPAI H7N9 virus GD/16 was kindly provided by Prof. Yoshihiro Kawaoka. The post-infection chicken antiserum raised against recombinant LPAI H7N9 AH/13 was generated by infection of specific-pathogen-free (SPF) Rhode Island Red chickens (Roslin Institute, UK) with the virus (28). The sera were heat-inactivated at 56°C for 30 min before further analysis.

### Plaque assay

The plaque assay was carried out as previously described (29), MDCK cells were infected with H7N9 AIVs 10-fold serial diluted for 1 h at 37°C. The cells were then washed once with phosphate-buffered saline (PBS) and replenished with Minimum Essential Medium (MEM)-agarose overlay. The MEM-agarose overlay medium contains 1× MEM, 0.6% agarose (Thermo Fisher Scientific), 100 units/mL penicillin, 100 µg/mL streptomycin, 2 mM L-glutamine, 0.3% bovine serum albumin (BSA), 15 mM 4-(2-hydroxyethyl)-1-piperazineethanesulfonic acid (HEPES), 0.22% sodium bicarbonate and 0.01% Diethylaminoethyl (DEAE)-Dextran (Sigma-Aldrich). The cells were further incubated at 37 °C for 48 h before being fixed with acetone and methanol (1/1 in volume) for 10 minutes (min). The plaques were visualized by incubating cells with mouse anti-NP monoclonal antibody (ATCC) (1 in 500 dilution) diluted in blocking buffer containing PBS with 5% FCS for 1 h at room temperature. Cells were subsequently rinsed with PBS supplemented with 0.01% Tween 20 and probed with horseradish peroxidase-labelled rabbit anti-mouse immunoglobulins (1 in 500 dilution) (DAKO, Agilent Technologies) for 40 min. After gentle rinsing with PBS supplemented with 0.01% Tween 20, cells were stained with 3,3′-diaminobenzidine (DAB) substrate-chromogen solution (DAKO) for 7 min. The plaque images were taken by Panasonic camera FZ38 and 30 plaques were randomly selected for each virus to measure the plaque diameter.

### *In ovo* growth of H7N9 AIVs

Ten-day-old embryonated chicken eggs were infected with 100 plaque-forming units of H7N9 AIVs. The eggs were incubated at 35°C for 48 h before allantoic fluid was harvested and titrated by HA assay.

### HA assay

The HA titration assays were performed with 1% chicken red blood cells as described in the WHO animal influenza training manual (30). Briefly, 50 μl of viruses were two-fold serially diluted in 96-well V-bottom microtitre plates (Greiner), followed by mixing with of 50 μl of 1% washed chicken red blood cells. The plates were incubated for 30 min at room temperature before the HA titre was recorded as the reciprocal dilution of the last well which contained non-agglutinated red blood cells and presented as HA units/50 μl.

### Hemagglutinin inhibition (HI) assay

25 μl of ferret sera or human mAbs were two-fold serially diluted in 96-well V-bottom plates (Greiner) with PBS prior to being mixed with equal volume of H7N9 Anhui/13 (8 HA units). The concentration for human mAb L4A-14, K9B-122, L3A-44, L4B-18 is 1.0 mg/ml, 1.5 mg/ml, 0.8 mg/ml, and 1.0 mg/ml respectively. The antibody-virus mixtures were incubated for 1 h at room temperature and then mixed with 50 μl of 1% chicken RBC. The plates were incubated for further 30 min at room temperature before the HI titer was recorded as the as log2 of the reciprocal of the dilution.

### HA thermal stability assay

The recombinant H7N9 AIVs were diluted with allantoic fluid harvested from non-infected embryonated chicken eggs to 64 HA units/50 μl. The viruses were then heated at 50°C, 50.7°C, 51.9°C, 53.8°C, 56.1°C, 58.0°C, 59.2°C and 60°C using PCR thermal cycler (Biorad) for 30 min or left at 4°C as control before HA assay.

### Syncytium formation assays

The pH of fusion for H7N9 AIV was determined by syncytium formation assays as previously described (9). To determine the optimal concentration of viruses to use in syncytium assay, Vero cells in 96-well-plate were infected with H7N9 AIVs two-fold serially diluted in DMEM for 1 h. The inoculum was then removed and washed with PBS before addition of DMEM medium with 10% FCS for 15 h. The cells were then fixed with methanol and acetone (1:1 in volume) mixture and then immuno-stained with an anti-nucleoprotein (NP) mouse mAb followed by horseradish peroxidase-labelled rabbit anti-mouse immunoglobulins (DAKO) as previously described (31). The virus concentration with the highest dilution still infecting 100% of the Vero cells was used to infect Vero cells in 96-well plate for 1 h. The inoculum was then removed and washed with PBS before the addition of DMEM medium with 10% FCS for 15 h. At 16 h post-infection, cells were treated with 3 μg/ml tosylsulfonyl phenylalanyl chloromethyl ketone (TPCK)-treated trypsin for 15 min and then exposed to PBS buffers with pH values ranging from 5.2 to 5.9 (at 0.1 unit increments) for 5 min. The PBS buffer was then replaced with DMEM with 10% FCS, and the plates were further incubated at 37 °C for 3 h to allow for syncytium formation. To visualize the syncytium formation, the cells were fixed with methanol and acetone (1:1 in volume) mixture and stained with Giemsa stain (Sigma-Aldrich) for 3 h at room temperature. The pH at which around 50% of maximum syncytium formation estimated was taken as the predicted pH of fusion. Images were taken on the Evos XL cell imaging system (Life Technologies).

### Virus purification and biolayer interferometry

H7N9 AIVs were purified as previously described (32). Briefly, virus harvested from allantoic fluid of embryonated eggs was pelleted by ultracentrifugation and then purified through a continuous 30 to 60% (weight/volume) sucrose gradient and resuspended in PBS buffer. The biotinylated α2,3- and α2,6-linked sialyl lactosamine sugars (3SLN and 6SLN, respectively) were purchased from GlycoNZ. Virus was diluted in HBS-EP buffer (TEKnova) containing 10 μM oseltamivir carboxylate (Roche) and 10 μM zanamivir (GSK) to a concentration of 100 pM, and the binding to receptor analogues was measured on an Octet RED instrument (ForteBio). The equilibrium responses for virus binding were plotted as a function of the amount of sugar immobilized on the biosensor calculated from the response during the sugar loading step (33). The relative dissociation constant, as a measure of binding to 3SLN, and 6SLN was calculated.

### Statistical analysis

Statistical analyses were performed using GraphPad Prism 8 (GraphPad Software). One-way ANOVA and Tukey’s multiple comparison test was used to analyse the differences between different groups. p values < 0.05 were considered significant.

## ACKNOWLEDGMENTS

The work described herein was funded by the UK Research and Innovation (UKRI), Biotechnology and Biological Sciences Research Council (BBSRC) grants: BB/X006166/1, UK-China-Philippines-Thailand Swine and Poultry Research Initiative (BB/R012679/1), Zoonoses and Emerging Livestock Systems (ZELS) (BB/L018853/1 and BB/S013792/1), the Global Challenges Research Fund (GCRF) One Health Poultry Hub (BB/S011269/1), the Responsive Mode (BB/R012679/1), Japan Partnering Award (BB/P016472/1), the Pirbright Institute strategic program grants (BBS/E/I/00007030, BBS/E/I/00007031, BBS/E/I/00007035 and BBS/E/I/00007036), the British Council Newton Fund Institutional Links grant (IL3261727271) and the Royal Society Newton Fund grant (NAF\R1\191166). The funders had no role in study design, data collection, data interpretation or the decision to submit the work for publication.

We thank Steve Martin for allowing us to use his Octet analysis software for the receptor binding analysis.

